# LncRNA *PAINT* is Associated with Aggressive Prostate Cancer and Dysregulation of Slug and Related Genes

**DOI:** 10.1101/2020.10.29.361105

**Authors:** Md Faqrul Hasan, Kavya Ganapathy, Jiao Sun, Khatib Ayman, Thomas Andl, Julia N. Saulakova, Domenico Coppola, Wei Zhang, Ratna Chakrabarti

**Affiliations:** Burnett School of Biomedical Sciences, University of Central Florida, Orlando, Florida; Department of Computer Science, University of Central Florida, Orlando, Florida; Moffitt Cancer Center, Tampa, Florida; Florida Digestive Health Specialists, Bradenton, Florida

**Keywords:** LNC-RNA, RNA-Seq, EMT, apoptosis, drug resistance

## Abstract

Long non-coding RNAs (lncRNAs) play regulatory roles in cellular processes and their aberrant expression may drive cancer progression. Here we report the function of a lncRNA *PAINT* (Prostate Cancer Associated Intergenic Non-Coding Transcript) in promoting prostate cancer (PCa) progression. Upregulation of *PAINT* was noted in advanced stage and metastatic PCa. Inhibition of *PAINT* decreased cell proliferation, S-phase progression, increased expression of apoptotic markers, and improved sensitivity to docetaxel and Aurora kinase inhibitor VX-680. Inhibition of *PAINT* decreased cell migration and reduced expression of Slug and Vimentin. Ectopic expression of *PAINT* suppressed E-cadherin, increased S-phase progression and cell migration. *PAINT* expression in PCa cells induced larger colony formation and higher expression of mesenchymal markers. Transcriptome analysis followed by qRT-PCR validation showed differentially expressed genes involved in epithelial mesenchymal transition (EMT), apoptosis and drug resistance in *PAINT*-expressing cells. Our study establishes an oncogenic function of *PAINT* in PCa.

## Introduction

Long non-coding RNAs (lncRNAs) have emerged as key regulatory molecules that play vital roles in gene regulation (1) and are frequently dysregulated in cancers. Due to their heterogeneity, LncRNAs have been integrated in many molecular processes including regulation of genomic integrity, cell fate decisions, differentiation, development, metabolism and cell death (2, 3). Therefore, it is conceivable that lncRNAs also play crucial roles in the initiation and progression of malignancies. Because of their association with tumorigenesis, lncRNAs are becoming novel candidates for therapeutic interventions.

Prostate cancer (PCa) is a commonly diagnosed cancer in American men. Functional vicissitudes of oncogenes and tumor suppressors involving multiple signaling pathways have been implicated in promoting metastatic and drug-resistant PCa (4). In this regard, a number of aberrantly expressed lncRNAs have significant impacts on the development, metastatic progression and emergence of drug-resistant PCa (5). For example, lncRNA HOTAIR and SCHLAP1 are commonly upregulated in advanced prostate cancer and promote drug resistance and aggressiveness (6, 7). However, lncRNA MEG3 and LincRNA-p21 are often downregulated in PCa and act as tumor suppressors (8). Although the functional involvement of a number of lncRNAs have been studied in PCa, a vast majority of dysregulated lncRNAs lack functional characterization and thus may play important roles in PCa progression.

Here we show functional characterization of a long non-coding RNA *PAINT* (Gene ID: LINC00888, Acc.# NR_038301.1) that promotes PCa progression. We demonstrated that *PAINT* is upregulated in PCa and exhibits a positive correlation with clinical stages of PCa. Our data suggest that *PAINT* promotes PCa phenotypes through upregulation of mesenchymal marker Slug and its target genes by a collective activation of Wnt/◻-catenin signaling cascade and genes involved in epithelial-mesenchymal transition (EMT). We also show that inhibition of *PAINT* has a beneficial effect on drug sensitivity of aggressive PCa cells. Our findings provide a novel insight on the role of lncRNA *PAINT* in progression of aggressive PCa.

## Materials and Methods

### Patient Tissues

PCa tissue microarray (TMA) with 63 cores of de-identified prostate tissues (US Biomax) composed of samples from normal prostate and prostate adenocarcinoma from stages I, II, III and IV were used for analysis of *PAINT* expression. TNM classification and Gleason Scores of samples were included as pathological criteria of the tumors (Supplemental Data 1 Table 1).

### RNA In-situ Hybridization

Formalin-fixed paraffin embedded (FFPE) TMA slides were used with *PAINT*-specific oligonucleotide probes (NR_038301.1) and positive control probes, which were designed and synthesized by ACD diagnostics and used for automated RNAScope LS assays compatible with Leica Biosystems’ BOND RX System. FFPE slides were pretreated and processed for probe hybridization, signal amplification through binding of alkaline phosphatase labeled probes and addition of Fast Red substrate for signal detection using RNAScope 2.5 LS reagents red kit (ACD) and manufacturer’s protocol (9). Individual images were scanned by AperioScope (Leica) and analyzed using QuPath software and expression of *PAINT* were counted as red dots/100 cells (10). Positive signals were also scored by a pathologist (D.C.) using the Allred scoring system (11).

### Cell Line Maintenance and Transfection

PC-3 cells (RRID:CVCL_0035; obtained from ATCC) were cultured in F-12 Kaighn’s Modification HAM medium (Sigma Aldrich) containing 10% heat-inactivated Fetal Bovine Serum (FBS) (Atlanta Biologicals) and 1% antibiotic/antimycotic (Life Technologies). C4-2B cells (RRID:CVCL_4784; obtained from ATCC) were maintained in RPMI-1640 media (Sigma Aldrich) containing 10% heat-inactivated FBS and 1% antibiotic/antimycotic. 22Rv1 cells (RRID:CVCL_1045; obtained from ATCC) were maintained in RPMI-1640 medium containing 10% heat-inactivated FBS and 1% antibiotic/antimycotic. Androgen dependent LNCaP subline LNCaP-104S cells (RRID:CVCL_M126; obtained as a gift from Dr. Shutsung Liao from University of Chicago) were maintained in DMEM media (Sigma Aldrich) containing 10% heat-inactivated FBS and 1% antibiotic/antimycotic and 1 ng/mL Dihydrotestosterone (DHT) (Life technologies). MDA-PCa-2b cells (RRID:CVCL_4748; obtained from ATCC) were maintained in 10% F-12K medium containing 10% non-heat inactivated FBS, 1% antibiotic/antimycotic, 25ng/mL cholera toxin, 10ng/mL mouse EGF, 0.005 mM phosphoethanolamine, 100 pg/mL hydrocortisone, 45nM sodium selenite, 0.005 mg/mL human. recombinant insulins. RWPE-1 cells (RRID:CVCL_3791; obtained from ATCC) were sub-cultured and maintained in Keratinocyte Serum Free Medium (K-SFM) (Gibco) supplemented with 0.05 mg/mL BPE and 5ng/mL EGF (provided with K-SFM kit). Both LNCaP C4-2B and LNCaP 104-S are derivatives of LNCaP (RRID:CVCL_0395). All cell lines except RWPE1 have been authenticated using STR profiling within the last three years and tested for mycoplasma contamination by DAPI staining. No STR profiling of RWPE1 cells was done as this cell line was obtained from ATCC immediately before conductng the experiments. All experiments were performed with mycoplasma-free cells.

PC-3 cells were transfected with *PAINT* siRNA smart pool PC-3-*PAINT*^si^ and non-targeting siRNA pool PC-3^C^ (negative control) (Dharmacon) using RNAiMAX (Invitrogen) for knock-down studies. Cells were harvested 48h or 72h after transfection for subsequent experiments. For overexpression studies, *PAINT* overexpressing C4-2B subline (C4-2B-*PAINT*^++^and control C4-2B (C4-2B^C^) subline were generated by transfecting C4-2B cells with either pcDNA3.1+*PAINT* or pcDNA3.1+ control using Lipofectamine 3000 (Invitrogen). Colonies were selected by treating transfected C4-2B cells with 1 mg/mL of G-418 (KSE Scientific) for 3 weeks and cloned for generating stable sublines. *PAINT* overexpression was determined using qRT-PCR for each subline and used for relevant experiments.

### Quantitative Real-Time PCR

Total RNA was extracted from different PCa cell lines using Direct-zol quick miniprep plus RNA extraction kit (Zymo Research). cDNA was synthesized from extracted RNA using RT^2^ First Strand Kit (Qiagen) or High-Capacity cDNA Reverse Transcription Kit (Applied Biosystems) as suppliers’ recommendation. Expression of *PAINT* was determined using *PAINT*-specific primer pairs (RT^2^ qPCR Primer Assays - Qiagen) and internal control EIF3D and RPL13A specific primers (RT^2^ qPCR Primer Assays -Qiagen) and RT² SYBR® Green qPCR master mix using the recommended protocol. Quantitative RT-PCR were performed in a QuantStudio 7 thermal cycler (Applied Biosystems) and was quantified based on SYBR green fluorescence and normalized based on the passive reference dye, ROX. Acquired data was analyzed based on 2^−ΔΔCT^ Livak-method and our published study (12) to identify expression of PAINT in relevant experiments.

### Western Blotting

Total protein lysates were prepared from PC-3-*PAINT*^si^, PC-3^C^ and C4-2B-*PAINT*^++^ and C4-2B^C^ sublines using RIPA buffer supplemented with phosphatase and protease inhibitors (Fisher Scientific) and used for immuno-blotting using anti-Slug, anti-Vimentin, anti-E-Cadherin, anti-PARP, anti-cleaved-Caspase 3, anti-Beta-Catenin, anti-phospho-AKT, pan-Akt, alpha-tubulin and anti-GAPDH (Cell Signaling Technology), and anti-PCNA (Santa Cruz). Alpha-Tubulin or GAPDH were used as internal controls. Blots were imaged using ECL chemiluminescence substrates and imaged with ChemiDoc MP Imaging System (Bio-Rad). Comparative expression was performed based on densitometry analysis using Image J software.

### Drug Sensitivity Assay

For drug sensitivity assay, transfected PC-3 cells were seeded in 96 well plates and transfected with siRNAs. At 24h after transfection, cells were treated with DTX at 5nM and 25nM or VX-680 at 1μM or DMSO as the control and continued incubation for additional 48h. MTS assays were performed to quantify viable cells at different experimental conditions

### Preparation of RNA Samples for Next Gen RNA-Sequencing

Total RNA was depleted of rRNAs using Arraystar rRNA removal kit and used for library preparation using Illumina kit for the RNA-seq library preparation. This includes RNA fragmentation, random hexamer priming for the first stand and dUTP based second strand synthesis followed by A tailing and adapter ligation. Next PCR amplification was performed for generating cDNAs for library preparation. The quality of the RNA library was checked for integrity of fragments between 400-600 bases Agilent 2100 Bioanalyzer and quantified using qRT-PCR through absolute quantification. DNA fragments were denatured using 0.1M NaOH and sequencing was performed in Illumina NovaSeq 6000 after fragments were amplified using NovaSeq 6000 S4 Reagent Kit for 150 cycles. RNA-Seq library preparation, sequencing and data analysis was performed by Arraystar Inc. The Raw data file in FASTQ format was subjected to quality control plot using FastQC software to obtain a quality score. All samples had a Q≥30 score of ≥93. Next, the fragments were adapter trimmed and filtered ≤20 bp reads using cutadapt software, and trimmed fragments aligned to reference genome (including mRNA, pre-mRNA, poly-A tailed lncRNA and pri-miRNA) with HiSAT2 (24) software. More than 92% of the reads of the trimmed pairs were aligned with the reference genome.

### RNA Sequencing and Data Analysis

Whole genome transcription profiling except ribosomal RNAs (rRNAs) and transfer RNAs (tRNAs) was performed using C4-2B-*PAINT*^++^ or C4-2B^C^ cells. Quantification of FPKM values and differentially expressed gene and transcript analyses were performed using R package Ballgown. Fold change (cutoff 1.5), *p-value* (≤0.05) and FPKM (≥ 0.5 mean in one group) were used for filtering differentially expressed genes and transcripts. GO enrichment analysis was used to associate the differentially expressed genes to specific GO terms. Pathway analysis using KEGG database was done for determining the enrichment of specific pathways by the differentially expressed genes. The *p-values* calculated by Fisher’s exact test was used to estimate the statistical significance of the enrichment of GO terms and pathways between the two groups. All other analysis and statistical computing were performed using R, Python and Shell environment by Arraystar Inc.

### Statistical Analysis

For TMA analysis, the key measures of interest (dependent variables) was the average number of red spots per 100 cells in a section computed over three sections from the same tumor tissue; the average was treated as a continuous variable. Table 4 presents the summary statistics for the three numbers (termed targets) and the average number. Additional measures (independent variables) for cancer tissues included cancer stage (I/II, III, IV), grade, Gleason Score Indicator (GSI) that differentiated between “low” (6 or less) and “high” (7 or greater), and metastasis indicator that differentiated between tissues with (TNM contained N1, N2, M1, M1b, or M1c) and without metastases (TNM contained none of N1, N2, M1, M1b, and M1c). We also used a multinomial regression model, Kaplan-Meier estimation, one-way ANOVA and two-sample t-tests. The significance levels were fixed at the 5% level (*p*-value≤0.05) or for some results at 10% level (*p*-value≤0.1). Multiple comparisons were performed using Bonferroni adjustments. Analyses were performed using SAS®9.4 software (12). Data were represented as mean ± SD.

Methodologies of all phenotypic experiments such as cell proliferation, flow cytometry and Annexin V apoptosis assays, migration and colony formation assays and immunofluorescence assays are provided in the supplemental information methods sections.

## Results

### *PAINT* is upregulated in aggressive PCa

To explore the expression of *PAINT* in PCa, we performed *PAINT* RNA-in-situ hybridization (RNA-ISH) using PCa TMA comprised of normal prostate tissues and prostate adenocarcinomas with pathological criteria of stages I/II, III and IV (Suppl. Data 1 Table 1). We noted overexpression of *PAINT* in prostate tumors compared to the normal prostate tissues specifically, in late stage PCa (stage III and stage IV) compared to early stage PCa (Figures 1A & 1B). Group sample sizes for the statistical analysis and the summary statistics for the primary measures for cancer and normal tissues is shown in supplementary data (Suppl. Data 2 Table 2 and 3).

**Figure 1:**
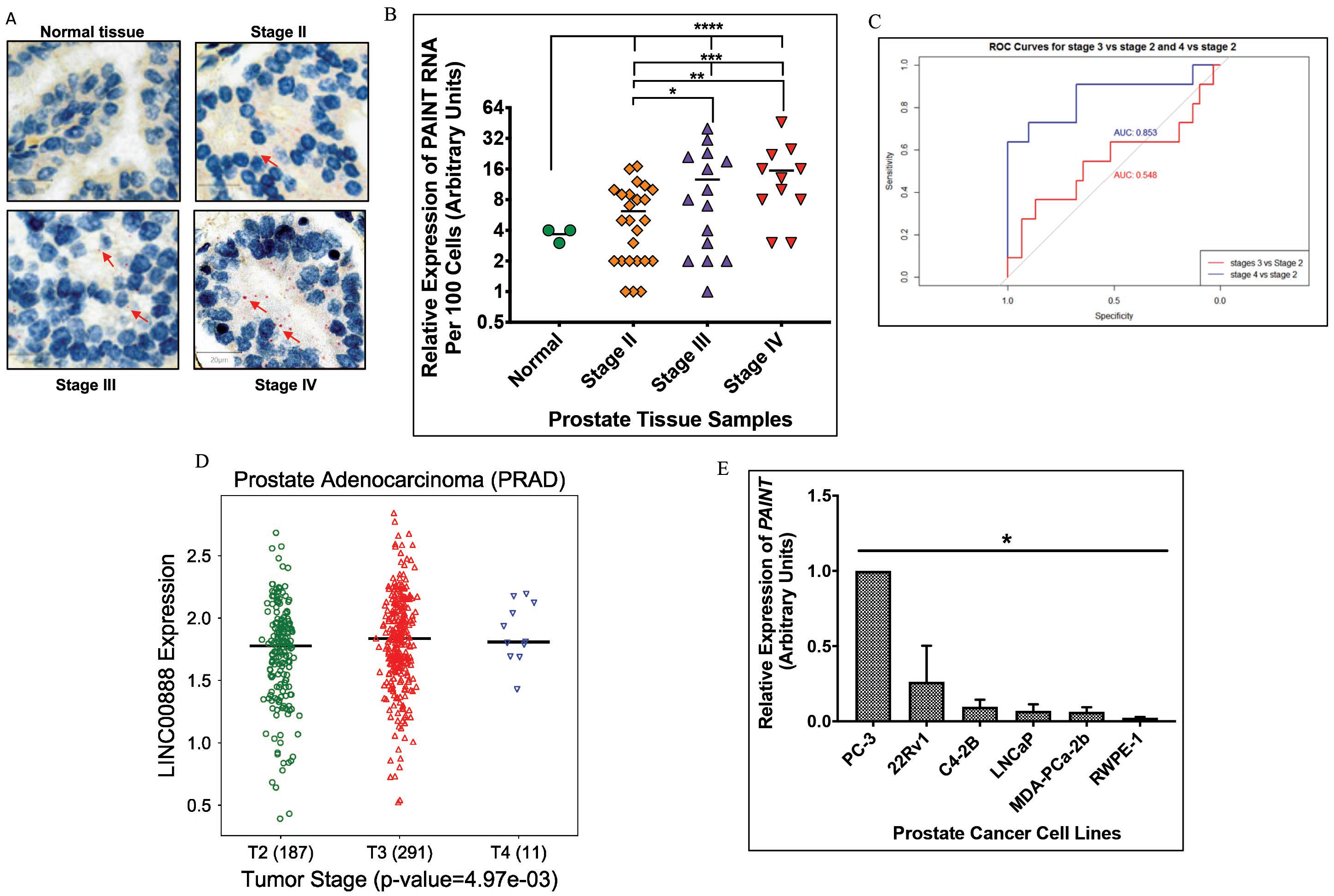
*PAINT* is upregulated in late stage prostate cancer. A: Representative TMA images of *PAINT* RNA-ISH in tissues from normal prostate stage II, stage III and stage IV PCa. Arrows: positive signals (Red dots). B: Comparative expression analysis of *PAINT* in prostate cancer and normal tissues \**p-value*=0.02, \*\**p-value*=0.0022, \*\*\**p-value*=0.013, \*\*\*\**p-value*=0.016. C: Receiver Operating Characteristic (ROC) curves based on the two models. Blue line illustrates that the area under the ROC curve is 0.86 confirming a high level of accuracy of predicting stage IV versus stage I/II. Red line illustrates a poor level of accuracy of predicting stage III versus stage I/II. D: Analysis of RNA-seq data using TCGA PRAD dataset shows higher expression of *PAINT* (LINC00888) in stage III and stage IV compared to stage II PCa tissues. E: Expression analysis showing highest expression of *PAINT* in metastatic PC-3 cells. Data represent mean ± SD of three biological replicates. * *p* < 0.0001.

The summary statistics showed that the model for *PAINT* expression was significant (*F*=342.6, *df*=7, *p-value*<0.001, *R^2^*=0.47) and indicated significance of stage (*p-value*<0.001) and metastasis (*p-value*<0.001) (Suppl. Data 3 Table 4). Analysis showed significant differences between stages I/II and IV (adjusted *p-value*<0.003) and stages III and IV (adjusted *p-value*=0.039). We note that grade was significant at 10% level (*p-value*=0.057). Analysis of the prediction of PCa stages incorporated a multinomial regression model (Likelihood Ratio *Χ^2^*=16.5, *df* =8, *p-value*=0.036, *R^2^*=0.27) for the logit of probability of stage IV versus stage I/II and probability of stage III versus stage I/II. Our results showed that *PAINT* expression was a significant predictor overall (*p-value*=0.010) and of the odds of cancer stage IV relative to stage I/II (*OR*=1.30, 95% *CI*=1.10:1.54, *p-value*=0.03) (Fig. 1C and Suppl. Data 4 Table 5). Analysis of TCGA PRAD (13) dataset further revealed higher expression of *PAINT* in late stage PCa tissues and was correlated to poor survival (Figure 1D, Suppl. Fig S1A). Analysis of *PAINT* expression in PCa cell lines showed its highest expression in metastatic PC-3 cells compared to other PCa and non-tumorigenic cell lines (Figure 1E). Collectively, these observations demonstrate that *PAINT* is upregulated in PCa tissues exhibiting a direct correlation with tumor stages and metastatic PCa.

### *PAINT* regulates cell phenotype and drug sensitivity in PCa cells

The functional role of *PAINT* in PCa was determined using knockdown and overexpression approaches. siRNA-based inhibition of *PAINT* in PC-3 cells (PC-3-*PAINT*^si^) and ectopic expression of *PAINT* in C4-2B cells (C4-2B-*PAINT*^++^) were used for subsequent studies. PC-3-*PAINT*^si^ cells showed an altered cell morphology from its spindle shape to a more epithelial cuboidal shape, reduced cell proliferation (26%) and S-phase cells (Fig. 2A and B and Suppl. Fig S1B, Figure 2E and F) compared to the control PC-3 (PC-3^C^) cells. Instead, C4-2B-*PAINT*^++^ cells showed increased cell proliferation (53%) (Figure 2C and D), higher Ki67 proliferation index (14) (Suppl. Fig S1C) and an enrichment in S-phase cell population (Figure 2G and H) compared to control C4-2B (C4-2BC) cells. Expression of S-phase marker PCNA (15) showed increased expression (40%) in C4-2B-*PAINT* ^++^ cells (Figure 2K and L) and reduced expression (18%) in PC-3-*PAINT*^si^ cells (Figure 2I and J) suggesting that *PAINT* expression influences cell cycle progression and cell proliferation.

**Figure 2:**
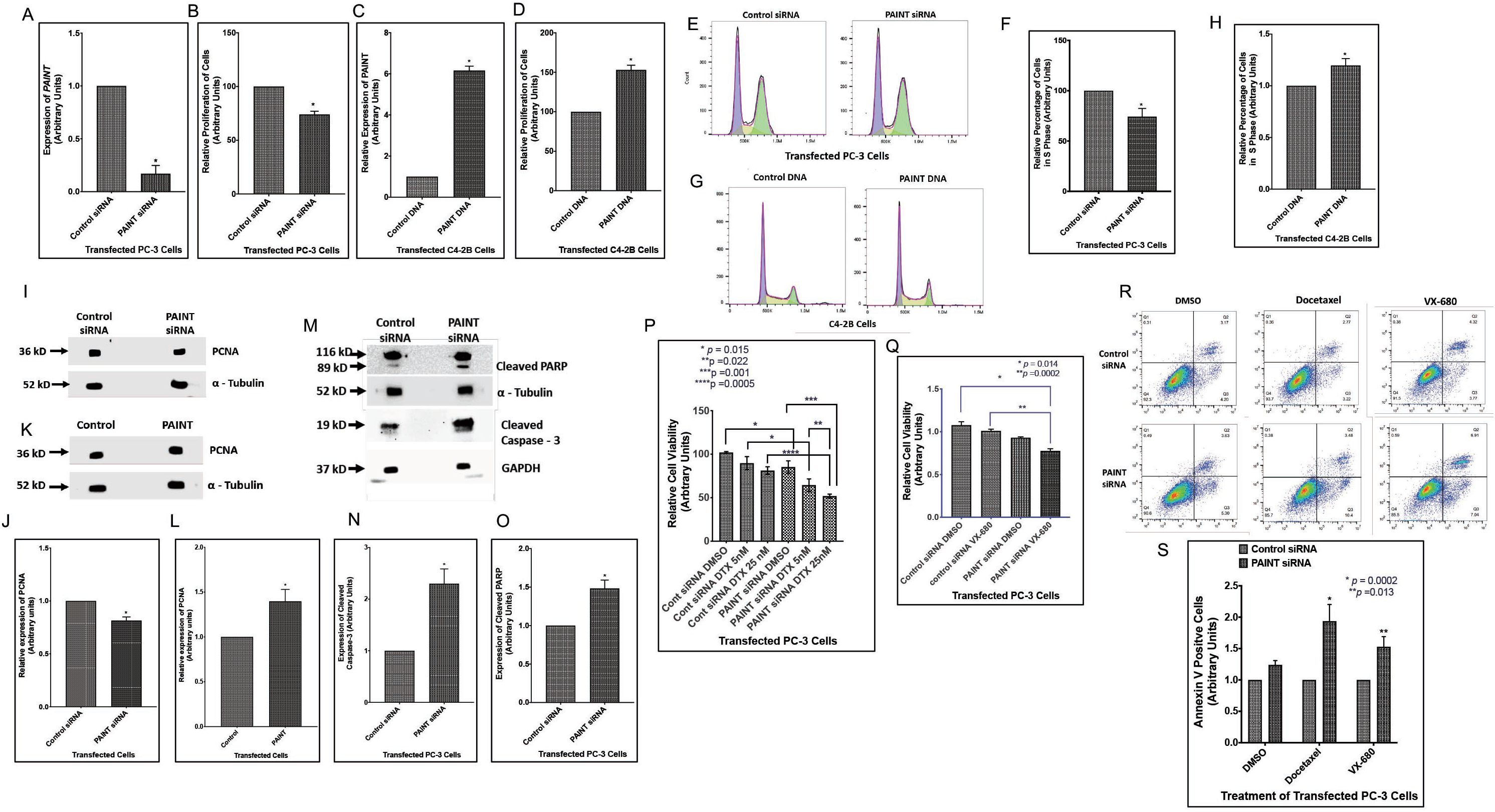
Changes in *PAINT* expression regulate cell proliferation, cell cycle progression, cell survival and drug sensitivity. A: Expression of *PAINT* in PC-3-*PAINT*^si^ cells. Data show the mean ± SD of three individual experiments. \**p-value* < 0.05. B: MTS assays showing proliferation of PC-3-*PAINT*^si^ and PC-3^C^ cells. C: qRT-PCR analysis showing expression of *PAINT* in C4-2B-*PAINT* ^++^ and C4-2B^C^ cells. Data represent the mean ± SD of three independent experiments. \**p-value*= 0.018. D: MTS assays showing proliferation of C4-2B-*PAINT*^++^ and C4-2B^C^ cells. Data represent the mean ± SD of three biological replicates. \**p-value* = 0.004. E: Two parameter histograms of propidium iodide (PI) stained PC-3^C^ (left) and PC-3-*PAINT*^si^ (right) cells. F: Comparative analysis of S phase cells showing a reduction in S phase population of PC-3-*PAINT*^si^ cells compared to PC-3^C^ cells. Data show the mean ± SD of 4 biological replicates. \**p-value* = 0.008. G: Representative cell cycle histogram of PI stained C4-2B^C^ (left) and C4-2B-*PAINT* ^++^ (right) cells. H: Comparative analysis of S phase cells showing an increase (19%) in S phase population of C4-2B-*PAINT* ^++^ cells compared to C4-2B^C^ subline. Data represent the mean ± SD for 3 biological replicates. \**p-value* = 0.036. I and K: Representative Western blot images of PCNA expression in PC-3-*PAINT*^si^ and C4-2B-*PAINT*^++^cells, respectively. Alpha tubulin was used as the loading control. J and L: Densitometric analysis of PCNA in PC-3-P *AINT*si and C34-2B-*PAINT*s++ with \**p-value =* 0.029 and \**p-value* = 0.024. respectively. M: Western blot images of PARP, activated PARP and cleaved Caspase-3 upon silencing of *PAINT* in PC-3-*PAINT*^si^ cells. N and O: Densitometric analysis of cleaved Caspase −3 and activated PARP in PC-3-*PAINT*^si^ cells compared to PC-3^C^ cells with \**p-value* = 0.045 and *\*\*p-value* = 0.016, respectively. Data show the mean ± SD of three individual experiments. P: Viability assays of PC-3-*PAINT*^si^ and PC-3^C^ cells in combination with DTX or DMSO treatments. Data show the mean ± SD of three individual experiments. \**p-value* =0.015, \*\**p-value*=0.022, \*\*\**p-value* = 0.001, \*\*\*\**p-value* = 0.0005. Q: Viability assay of PC-3-*PAINT*^si^ cells to VX-680 treatments compared to DMSO and PC-3^C^ cells. Data show the mean ± SD of three individual experiments. \**p-value* =0.014, ** *p-value* = 0.0002. R: Representative scatter plots of PC-3^C^ (top panel) and PC-3-*PAINT*^si^ (bottom panel) cells treated with DMSO (first panel), DTX (second panel) and VX-680 (third panel) showing more cells in the third and fourth quadrants in PC-3-*PAINT*^si^ cells upon DTX and VX680 treatments compared to DMSO controls. S: Enumeration of Annexin V positive cells upon DTX (5nM) and VX-680 (25nM) treatment of PC-3-*PAINT*^si^ cells compared to PC-3^C^ cells.

Next, analysis of cell survival showed increased expression of cleaved-Caspase 3 (1.5-fold) and cleaved PARP (2.5-fold) in PC-3-*PAINT*^si^ compared to PC-3^C^ cells (Figure 2M, N and O). This led us to examine the effect of *PAINT* inhibition on drug sensitivity of PC-3 cells to docetaxel (DTX) and VX680 (Aurora kinase inhibitor) (16, 17). DTX and VX680 treatment showed a synergistic effect with *PAINT* inhibition, on reduced cell viability at ~20% and 10% levels, respectively compared to control (Figure 2P and Q). Annexin-V apoptosis assays showed a significant increase in the percentage of apoptotic cells upon treatment with DTX and VX-680 in PC-3-*PAINT*^si^ cells compared to PC-3^C^ cells (Figure 2R and S). These results suggest that *PAINT* supports cell survival by evading apoptosis and decreasing the efficacy of therapeutic agents.

### *PAINT* promotes tumorigenic and migratory behavior of PCa cells

Next, we examined the effect of *PAINT* overexpression on anchorage independent colony formation and migration properties. We noted a higher percentage (52%) of large colonies (>70 μm) in C4-2B-*PAINT*^++^ cells compared to C4-2B^C^ cells (19%) (Figure 3A and B). Furthermore, a 34% increased rate of migration of C4-2B-*PAINT*^++^ cells compared to C4-2B^C^ cells (Figure 3E and F) was noted, whereas inhibition of *PAINT* expression showed an opposite effect (Figure 3C and D).

**Figure 3:**
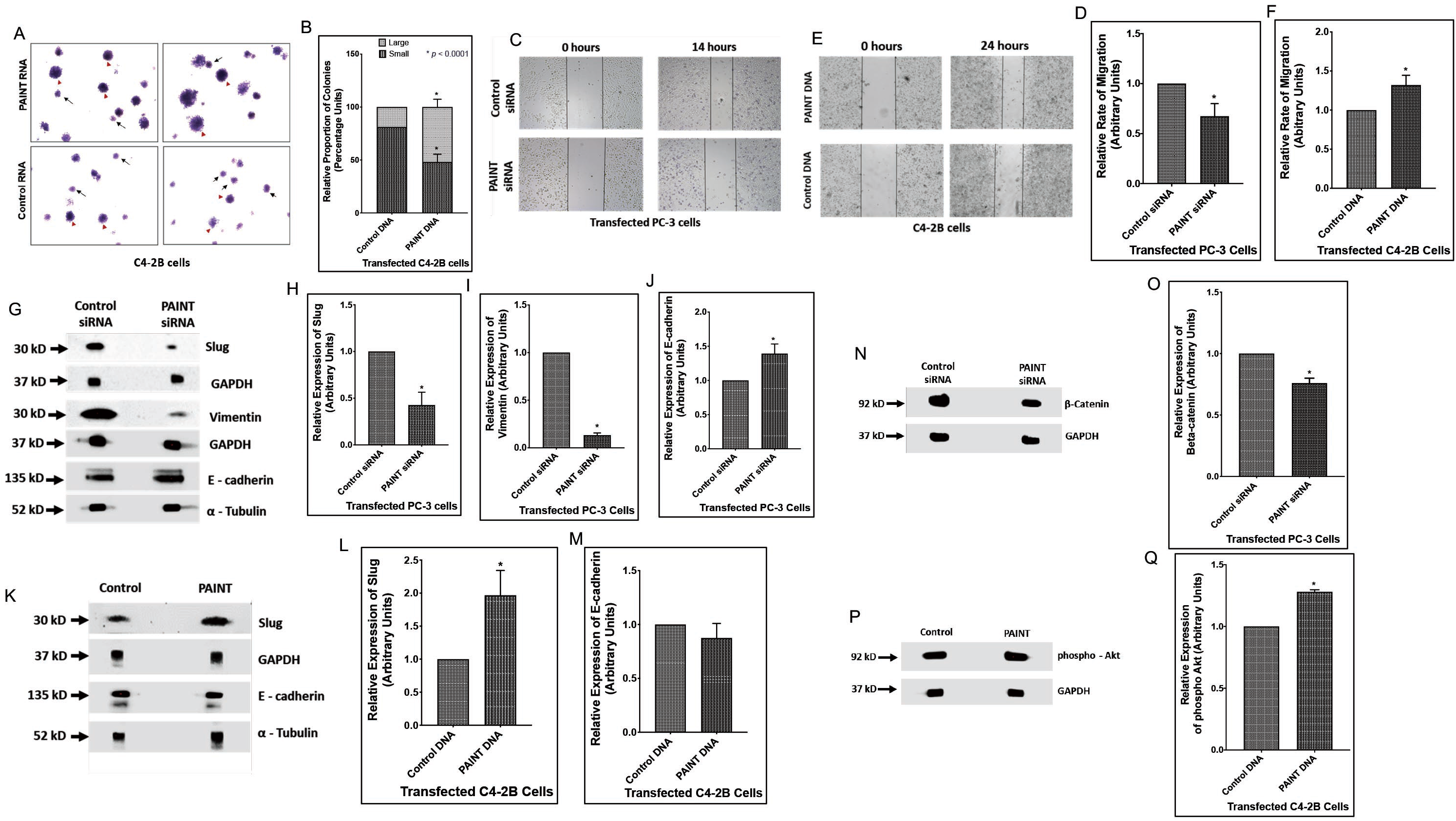
*PAINT* promotes colony formation, migration and epithelial-mesenchymal transition through modulation of multiple proteins. A: Representative images of soft agar colonies formed by C4-2B-*PAINT*^++^(top panels) and C4-2B^C^ cells (bottom panels). Red arrows represent large colonies and small black arrows represent small colonies. B: Quantitative analysis of the small (>7 μm) and large (>7 μm) of colonies formed by C4-2B-*PAINT*^++^ cells compared to C4-2B^C^ cells. Data show the mean ± SD of 3 biological replicates. \**p-value* < 0.0001. C: Representative images of migration of PC-3-*PAINT*^si^ cells compared to PC-3^C^ cells at 0 hours (left) and 14 hours (rights) after making the scratch. D: Analysis of the rate of migration of PC-3-*PAINT*^si^ cells compared to PC-3^C^ cells. Data show the mean ± SD of three independent experiments. \**p* = 0.046. E: Representative images of migration of C4-2B-*PAINT*^++^ cells (top) and C4-2B^C^ cells (bottom) at 0 hours (left) and 24 hours (right) after making the scratch. F: Analysis of the rate of migration of C4-2B-*PAINT*^++^ cells compared to C4-2B^C^ cells. Data represent mean ± SD for 3 biological replicates. \**p-value* = 0.01. G: Western blots showing altered expression of Slug, Vimentin and E-cadherin in PC-3-*PAINT*^si^ and PC-3^C^ cell lysates. GAPDH and ◻-tubulin were used as the loading controls. (H-J) Densitometry of Slug, Vimentin and E-cadherin expression in in PC-3-*PAINT*^si^ and PC-3^C^ cells. Data show the mean ± SD of three biological replicates. \**p-value* = 0.018, \*\**p-value* = 0.0002, \*\*\**p-value* = 0.011. K: Western blots showing expression of Slug and E-cadherin in C4-2B-*PAINT*^++^ and C4-2B^C^ cell lysates. GAPDH and ◻-tubulin were used as the loading controls. M: Densitometry of Slug and E-cadherin expression in C4-2B-*PAINT*^++^ or C4-2B^C^ cells. Data show the mean ± SD of three biological replicates. \**p-value* = 0.047. N and P: Western blots of ◻-catenin upon knockdown of *PAINT* in PC-3 cells, and GAPDH as the loading control. O: Densitometric analysis of ◻-catenin expression upon *PAINT* inhibition. Data represent mean ± SD. \**p-value* = 0.009. P: Western blots of phospho-Akt upon overexpression of *PAINT* in C4-2B cells and GAPDH as the loading control. Q: Densitometric analysis of phospho-Akt expression in C4-2B-*PAINT*^++^ subline compared to C4-2B^C^ cells. Data represent mean ± SD. \**p-value* = 0.0013.

Since EMT is frequently associated with metastatic and aggressive behavior, we focused on the relationship between *PAINT* and the key mesenchymal marker Slug (18, 19). Inhibition of *PAINT* expression in PC-3-*PAINT*^si^ reduced Slug by 57% compared to PC-3^C^cells. Regarding Slug-target genes, *PAINT* inhibition reduced Vimentin expression (87%), a Slug-induced gene (19), and increased E-Cadherin expression (40%), a Slug-repressed gene (20) (Figure 3G-J). Overexpression of *PAINT* in C4-2B-*PAINT*^++^reversed these effects showing an increase (96%) in Slug expression and a decrease (30%) in E-cadherin expression compared to C4-2B^C^ cells (Figure 3K-M).

As induction of EMT is often associated with activation of different signaling pathways, we monitored ◻-catenin expression which promotes EMT through Slug expression (21). A significant downregulation of ◻-catenin was noted in in PC-3-*PAINT*^si^ cells (Figure 3N and O). We also determined Akt activation, which can induce EMT and metastasis through Slug regulation (22), upon *PAINT* overexpression. Our results showed an increased expression of phospho-AKT while Akt levels remained unchanged compared to C4-2B^C^ cells (Figure 3P and Q). These observations indicate a role of *PAINT* in activating multiple signaling pathways that promote PCa progression and EMT.

### Transcriptome analysis revealed altered gene expression in PCa cells expressing *PAINT*

To understand the cellular reprograming behind the potential oncogenic role of *PAINT* in PCa progression, we performed RNA-seq analysis of C4-2B-*PAINT*^++^ (group E) and C4-2B^C^ cells (group C). The short reads were aligned to the human GRCh37 reference genome by HiSAT2 (23). Sequencing statistics of each sample is presented in Table 6 (Suppl. Data 5).

The abundance of genes and transcripts represented by FPKM values in the two group of samples were estimated by StringTie (24). Pearson Correlation analysis showed a strong correlation (>0.997) among *PAINT* overexpressing cells and control cells (Suppl. Fig. S2A). Using R package ballgown (25), a total of 76 upregulated genes and 61 downregulated genes with fold-change > 1.5 and *p-value* < 0.05 were identified in *PAINT*-expressing cells compared to control cells. A volcano plot based on log2 of fold change vs. -log10 of *p*-values of the genes showed a large magnitude of statistically significant changes between C4-2B *PAINT*-expressing cells compared to control cells (Figure 4A). Chromosomal mapping of dysregulated genes indicates that chromosomes 1, 11 and 19 contain the majority of the upregulated genes and chromosomes 1, 2 and 6 contain the majority of the downregulated genes (Fig, 4B). Unsupervised hierarchical clustering grouped the majority of differentially expressed genes based on their FPKM values (Figure 4C) showing distinct sets of genes that are upregulated or down regulated in *PAINT* expressing cells. Principal Component Analysis (PCA) shows distinct clustering of samples with genes that have p-value < 0.05 on FPKM abundance estimation (Figure 4D). In addition, 9,086 novel genes were identified using CPAT (Coding Potential Accessing Tool) (26), which showed distinct clusters of potentially protein coding and noncoding genes (Suppl. Fig S2B). Altogether, transcriptome analysis revealed a set of genes that were altered upon overexpression of *PAINT*.

**Figure 4:**
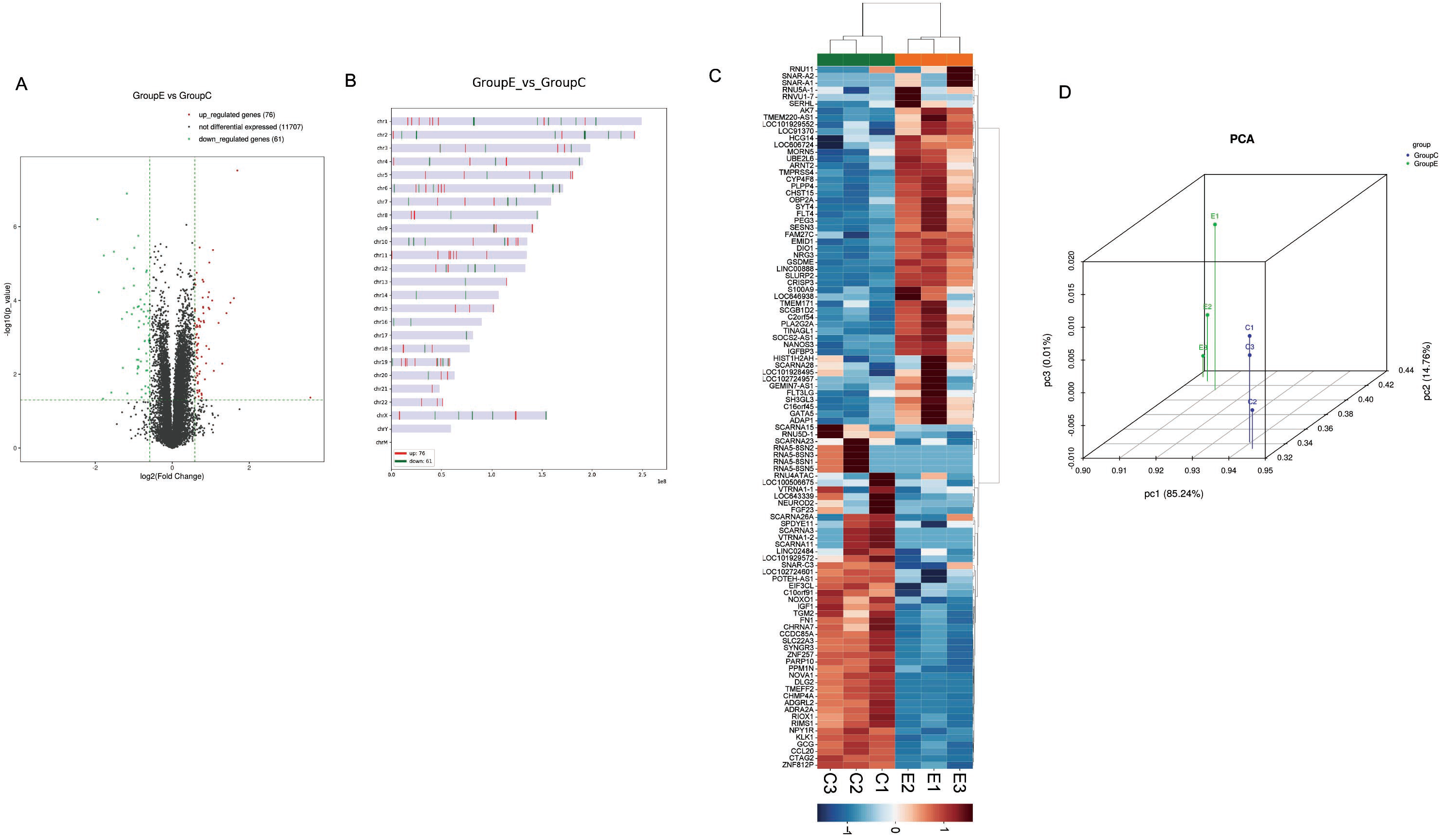
Transcriptome analysis reveals significantly dysregulated genes in *PAINT* overexpressing C4-2B cells. A: Volcano plot showing differential expression of genes with log2 transformed fold change gene expression (x-axis) and -log10 transformed *p*-*values* (Y-axis) between the two groups. Significantly upregulated and downregulated genes (*p*-*value* < 0.05 and fold change no less than 1.5) are marked with red and green dots respectively. Grey dots show genes that are not differentially expressed. B: Chromosomal maps showing location of differentially expressed genes on each chromosome in C4-2B-*PAINT*^++^ cells. C: Hierarchical Clustering analysis showed a significant number of differentially expressed genes between C4-2B-*PAINT*^++^ (E group) and C4-2B^C^ cells (C group). Genes are represented by rows and samples are represented by columns. Red color indicates higher expression and green color indicates lower expression. D: PCA of three biological replicates of C4-2B-*PAINT*^++^ cells compared to C4-2B^C^ cells showed distinguishable gene expression profiles between the two groups.

### Identification of functionally related groups and enrichment of pathways of dysregulated genes in *PAINT*-expressing cells

Gene Ontology (GO) analysis of top dysregulated genes was performed based on specific gene attributes such as Biological Process (BP), Molecular Function (MF) and Cellular Component (CC). Circular plots show the GO enrichment of the top downregulated genes (Suppl. Fig. S2C and D and Figure 5A) and upregulated genes (Suppl. Fig S2E and F, Figure 5C) in C4-2B-*PAINT*^++^ cells based on BP, CC and MF respectively. Furthermore, significantly downregulated (Figure 5B) and upregulated genes (Figure 5D) were grouped based on the top 10 enriched GO terms within BP, CC and MF. Furthermore, KEGG Pathway analysis of upregulated and downregulated genes revealed multiple pathways that were significantly enriched (Figure 5E and F, Suppl. Fig S3A-D, Suppl. Fig S3E-G). Taken together, gene ontology and KEGG pathway analysis establish that *PAINT* expression is associated with regulation of gene expression involved in several biological processes, functions and pathways which possibly contribute to prostate cancer progression.

**Figure 5:**
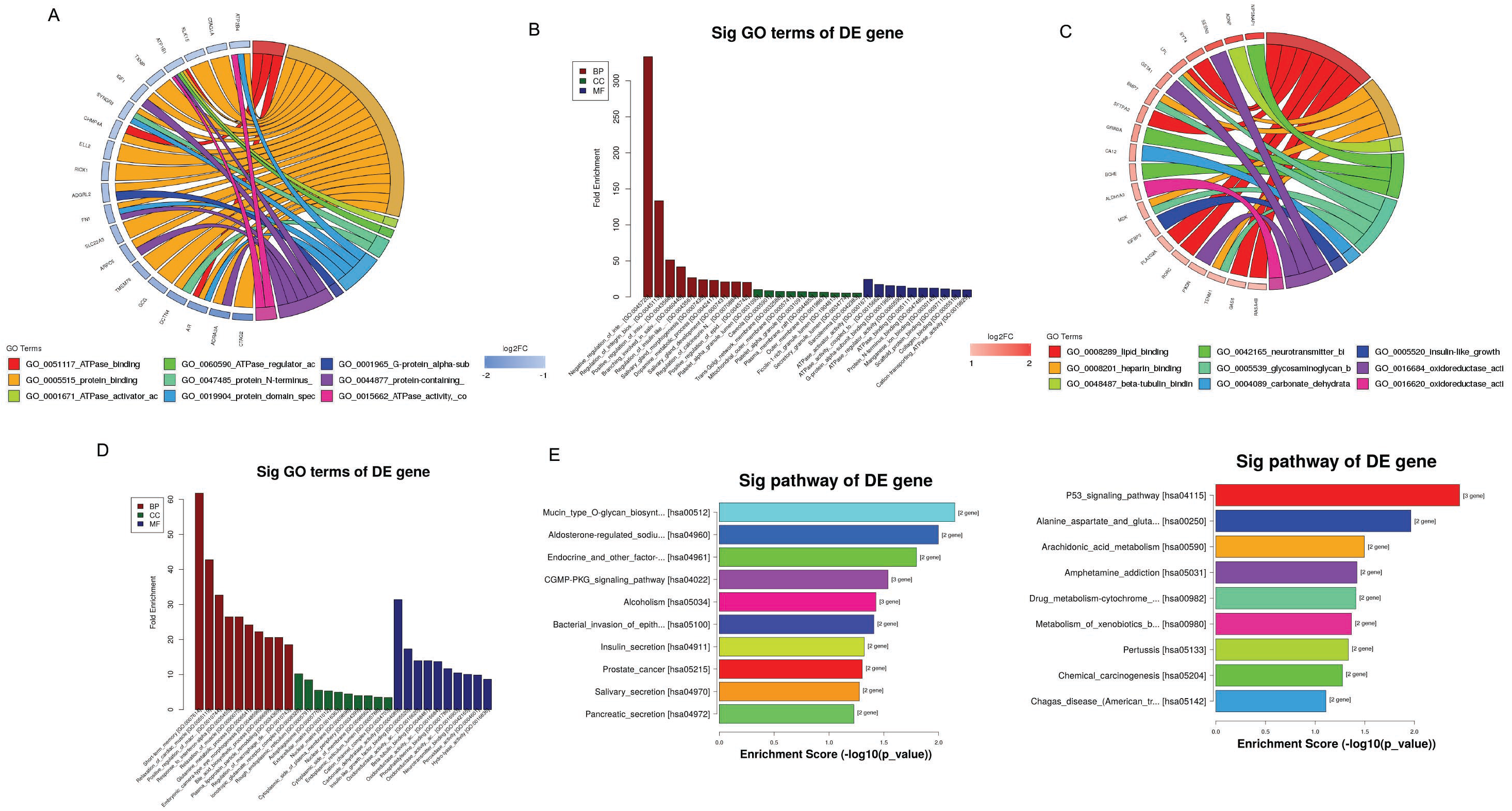
GO Enrichment and KEGG pathway analysis of differentially expressed genes between *PAINT* expressing C4-2B cells and control cells. A and C: Circular plots of GO enrichment analysis showing 20 downregulated genes and 20 upregulated genes in C4-2B-*PAINT*^++^ cells and their molecular functions, respectively. B and D: Top ten enriched GO terms for significantly downregulated genes and upregulated genes in C4-2B-*PAINT*^++^ cells based on biological process (BP), cellular component (CC) and molecular function (MF), respectively. The order of the bars is based on *p*-*value* from left to right (-log10). E and F: KEGG Pathway analysis showing top 10 pathways of downregulated genes and top 10 pathways for upregulated based on -log10(*p*-*value*) enrichment score.

### *PAINT*-expressing C4-2B cells reveal altered expression of gene targets involved in the EMT and apoptosis network that may regulate *PAINT*-induced PCa progression

Further analysis of the RNA-Seq data revealed a set of significantly dysregulated genes in *PAINT* expressing cells that were involved in EMT, apoptosis and drug sensitivity processes, similar to our observations from *in vitro* characterization studies (Figure 6A and B). The clinical significance of these genes in PCa progression was examined next using TCGA PRAD dataset (n=623). Our analysis identified two downregulated genes, TMEFF2 (27) (Fig 6C) and SLC22A3 (28) (Fig 6D) that showed decreased expression with stage-specific progression of PCa and five upregulated genes, TMPRSS4 (29) (Fig 6E), SYT4 (30) (Fig 6F), SESN3 (31) (Fig 6G), CRISP3 (32) (Fig 6H) and NANOS3 (33) (Fig 6I) that showed increased expression associated with stage specific progression of PCa. qRT-PCR analysis validated overexpression of selected five upregulated genes (Fig. 6J) and reduced expression of two selected downregulated genes (Fig. 6K). Altogether, our findings suggest that *PAINT* regulates a group of genes involved in cell growth, drug resistance and EMT, all of which comes together to drive PCa progression to more aggressive stages.

**Figure 6:**
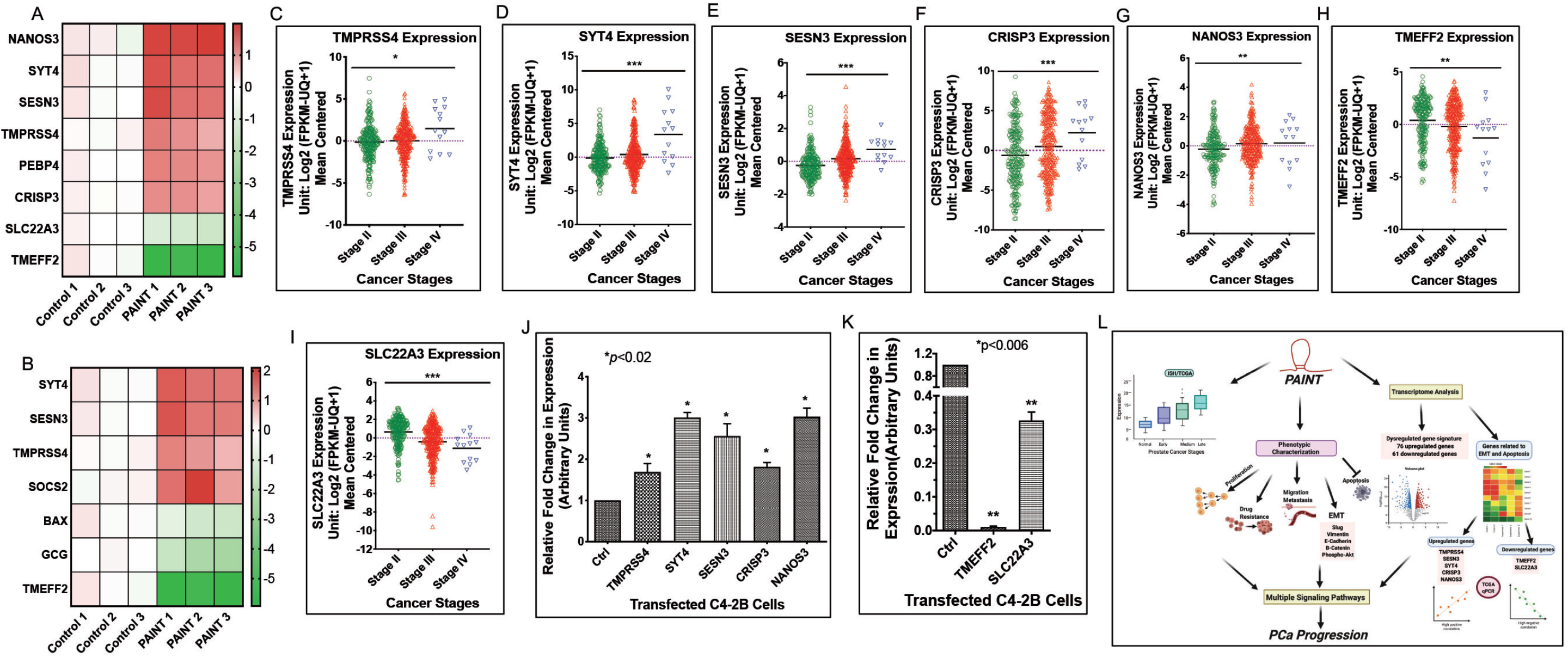
Functional analysis and validation of dysregulated genes in *PAINT* overexpressing prostate cancer cells. A and B: Expression heat map showing differential expression (Log2-fold) of genes involved in apoptosis (A) and EMT-regulated genes (B) between individual samples of C4-2B-*PAINT*^++^ (right) and C4-2B^C^ cells (left). Three biological replicates were included for each group. \**p-value* <0.05. (C-I) Expression of apoptosis and EMT related genes showing significant differences between stage II (n = 218) and stage III (313) and IV (n = 13) based on TCGA PRAD gene set analysis: TMPRSS4 (*p* = 0.02697) (C), SYT4 (*p* = 0.0041) (D), SESN3 (*p-value*< 0.0001) (E), CRISP3 (*p-value*=0.0006) (F), NANOS3 (*p-value*=0037) (G) TMEFF2 (*p-value* = 0.0018) (H) and SLC22A3 (*p*-value< 0.0001) (I). J: qRT-PCR validation of the selected five upregulated genes, TMPRSS4, SYT4, SESN3, CRISP3 and NANOS3 in C4-2B-*PAINT*^++^ and C4-2B^C^ cells. Data show the mean ± SD of three biological replicates. \**p-value* < 0.05. K: qRT-PCR validation of the selected two downregulated genes, TMEFF2 and SLC22A3 in C4-2B-*PAINT*^++^ and C4-2B^C^ cells. Data show the mean ± SD of three biological replicates. \**p*-*value* < 0.05. L: Functional overview of *PAINT* in prostate cancer progression.

## Discussion

Emerging studies established the role of aberrantly expressed lncRNAs in several cancers including PCa (34). Here, we focused on describing the function of a novel lncRNA and its role in PCa progression through modulation of specific gene networks. Our previous studies identified a tumor suppressor microRNA cluster, miR-17-92a, that is downregulated in PCa (35). RNA-seq analysis of PC-3 cells with restored miR-17-92a cluster miRNAs showed altered expression of several intergenic lncRNAs having more than a 2-fold change (log2) in expression, out of which *PAINT* was the most downregulated lncRNA. Expression of this lncRNA is upregulated in melanoma (36) but no information on the involvement of *PAINT* in PCa progression is available. Hence, we chose *PAINT* for further study on its role in PCa progression. To our knowledge, this is the first study that shows an oncogenic function of *PAINT* in PCa.

Our study revealed *PAINT* overexpression in prostate tumors and in metastatic PCa. TCGA data analysis further showed a positive correlation of *PAINT* with advanced stages of PCa with a poor patient survival. Similarly, *PAINT* was upregulated in metastatic and drug-resistant PCa cell line compared to the less aggressive and the nontumorigenic prostate cells. Collectively, *PAINT* upregulation in PCa, especially in the later stages, suggested the possibility of *PAINT* being a driver oncogene promoting progression of metastatic PCa.

The oncogenic function of *PAINT* was supported by the results showing *PAINT-*induced increased proliferation, migration, larger colony formation and EMT marker expression while most of these effects were blunted upon *PAINT-siRNA expression*. *PAINT* overexpression also facilitates cell survival as inhibition of *PAINT* induced apoptosis through activation of proteins involved in the apoptotic pathways (37) and as result, significantly improved the sensitivity of drug-resistant PC-3 cells to chemotherapeutic agents. These results highlight the importance of *PAINT* as a potential therapeutic target for PCa management.

Another hallmark of aggressive cancer is increased cell migration which contributes to the metastatic potential of cancer cells (38). Our results showed *PAINT’*s involvement in cell migration and a positive correlation of *PAINT* with Slug expression. Slug, one of the major transcription factors involved in EMT (39), promotes cell migration and invasion through modulation of different signaling pathways (40). As we noted, upregulation of Slug was associated with an increased expression of Vimentin, a Slug effector and a decreased expression of E-Cadherin that is negatively regulated by Slug (41, 42). Our results showing reduced *1*-Catenin expression in PC-3-*PAINT*^si^ cells and increased phospho-Akt in C4-2B-*PAINT*^++^ cells further suggest the involvement of Wnt signaling (43) and PI3K/Akt (44) signaling circuitries that affect Slug expression. Beta-Catenin, a key Wnt signaling pathway protein, regulates Slug expression and EMT transition via Vimentin and E-cadherin (39), while phospho-Akt regulates Slug expression via PI3K/Akt pathway (22). Both Wnt signaling (43) and PI3K/Akt (44) pathways are constitutively activated in PCa promoting cancer progression, metastasis and drug resistance. The cross-talk between signaling pathways further established the role of *PAINT* in EMT and activation of multiple signaling cascades that contributes to PCa metastasis.

Transcriptome profiling further complements the phenotypic characterization data of *PAINT* and provided a comprehensive understanding of genes involved in promoting PCa progression upon *PAINT* dysregulation. GO enrichment and KEGG pathway analysis revealed altered expression of genes and pathways indicating that *PAINT* may promote an oncogenic environment by simultaneously regulating various processes in PCa. Consistent with our observations from the *PAINT* expression associated phenotypic changes, transcriptome profiling identified a specific set of dysregulated genes involved in EMT, apoptosis and drug resistance. TCGA PRAD dataset analysis corroborated with our RNA-Seq data showing overexpression of TMPRSS4, SYT4, SESN3, CRISP3 and NANOS3 and reduced expression of TMEFF2 and SLC22A3 in PCa, which were further validated by qRT-PCR.

TMPRSS4 is overexpressed in PCa and other cancers (45), and is involved in EMT induction, specifically through modulation of Slug expression (29) and drug resistance (46). SYT4 is a neuroendocrine marker that is overexpressed during transition from localized to metastatic PCa (30) and in drug-resistant LNCaP cells (30). SESN3 (Sestrin 3) is implicated in promoting EMT (47) and inhibiting apoptosis in PCa (48). Inhibition of SESN3 increased sensitivity of drug-resistant PCa to cabazitaxel (31). CRISP3 and NANOS3 are highly upregulated in multiple cancers and promote EMT, migration and invasion (32, 33). TMEFF2 functions as a strong tumor suppressor by suppressing migration and invasion in PCa cells (49), and overexpression of TMEFF2 induced apoptosis in PCa (50). SLC22A3 is also downregulated in aggressive PCa (28) and functions as a direct inhibitor of EMT in esophageal cancer (50). These findings provide convincing evidence that *PAINT* plays an oncogenic role through modulation of different signaling molecules specifically involved in EMT, apoptosis and drug resistance, which collectively play an integrated role in PCa progression to a more aggressive and metastatic stage.

In summary, our findings establish *PAINT* as an oncogene in PCa and indicate the clinical significance of *PAINT* as a diagnostic marker and a possible therapeutic target for aggressive PCa (Fig. 6L). However, in-depth mechanism of *PAINT* mediated regulation of these cellular events, which promote PCa progression and metastasis is still unclear. Our future studies will focus on the mechanistic role of *PAINT* in functional regulation of different target genes and their involvement in progression of aggressive disease.

## Abbreviations

BP: Biological Process
C4-2B^C^: Control C4-2B cells without ectopic expression of *PAINT*
C4-2B-*PAINT*
^++^: C4-2B cells ectopic expressing *PAINT*
CC: Cellular Component
CPAT: Coding Potential Accessing Tool
DTX: Docetaxel
EMT: Epithelial-mesenchymal transition
FFPE: Formalin-fixed paraffin embedded
GO: Gene Ontology
GSI: Gleason score indicator
LncRNAs: Long noncoding RNAs
MF: Molecular Function
PAINT: Prostate Cancer Associated Intergenic Non-Coding Trancript
PCa: Prostate Cancer
PC-3-*PAINT*
^si^: siRNA-based inhibition of *PAINT* in PC-3 cells
PC-3^C^: Non-targeting siRNA pool transfected in PC-3
RNA-ISH: RNA in-situ hybridization
ROC: Receiving Operator Characteristics
rRNAs: Ribosomal RNAs
SD: Standard Deviation
TCGA PRAD: The Cancer Genome Atlas Prostate Adenocarcinoma Dataset
TMA: Tissue microarray
tRNAs: Transfer RNAs
VX680: Aurora kinase inhibitor

## Data Availability Statement

The RNA-Seq data generated in this study is available in GEO under accession number # GSE158953. All other data that support the findings of this study are available from the corresponding author upon reasonable request.

## Ethics Statement

All human tissues obtained from US Biomax as Tissue Microarrays (TMAs) were collected under HIPPA approved protocols using highest ethical standards with the donor being informed completely and with their consent.

## Acknowledgements

We thank Dr. Shaojie Zhang, Department of Computer Science, University of Central Florida, for his valuable comments and help in experimental design. This study is supported by a grant from NIH R21 (CA226611-01A1) to RC.

## Author Contributions

R.C., Md.H. and K.G designed the study. Md.H., K.G. and A.K. contributed to carry out the experiments. Md.H., K.G., A.K. and T.A contributed to analyzing the data. Md.H. contributed to analyzing the TMA staining and RNA-Seq data. D.C. performed the scoring of TMA and J.S performed statistical analysis of the data. W.Z and J.S. contributed to analyzing RNA-Seq data and TCGA PRAD data analysis. Md.H, K.G. W.Z and R.C. wrote the manuscript. R.C. and T.A. supervised the research. All authors read and approved the final manuscript.

## References

1. Ransohoff JD, Wei Y, Khavari PA. The functions and unique features of long intergenic non-coding RNA. Nat Rev Mol Cell Biol. 2018;19(3):143–57.

2. Fatica A, Bozzoni I. Long non-coding RNAs: new players in cell differentiation and development. Nat Rev Genet. 2014;15(1):7–21.

3. Sun Q, Hao Q, Prasanth KV. Nuclear Long Noncoding RNAs: Key Regulators of Gene Expression. Trends Genet. 2018;34(2):142–57.

4. Semenas J, Allegrucci C, Boorjian SA, Mongan NP, Persson JL. Overcoming drug resistance and treating advanced prostate cancer. Curr Drug Targets. 2012;13(10):1308–23.

5. Evans JR, Feng FY, Chinnaiyan AM. The bright side of dark matter: lncRNAs in cancer. J Clin Invest. 2016;126(8):2775–82.

6. Zhang A, Zhao JC, Kim J, Fong KW, Yang YA, Chakravarti D, et al. LncRNA HOTAIR Enhances the Androgen-Receptor-Mediated Transcriptional Program and Drives Castration-Resistant Prostate Cancer. Cell Rep. 2015;13(1):209–21.

7. Prensner JR, Iyer MK, Sahu A, Asangani IA, Cao Q, Patel L, et al. The long noncoding RNA SChLAP1 promotes aggressive prostate cancer and antagonizes the SWI/SNF complex. Nat Genet. 2013;45(11):1392–8.

8. Luo G, Wang M, Wu X, Tao D, Xiao X, Wang L, et al. Long Non-Coding RNA MEG3 Inhibits Cell Proliferation and Induces Apoptosis in Prostate Cancer. Cell Physiol Biochem. 2015;37(6):2209–20.

9. Wang F, Flanagan J, Su N, Wang LC, Bui S, Nielson A, et al. RNAscope: a novel in situ RNA analysis platform for formalin-fixed, paraffin-embedded tissues. J Mol Diagn. 2012;14(1):22–9.

10. Bankhead P, Loughrey MB, Fernandez JA, Dombrowski Y, McArt DG, Dunne PD, et al. QuPath: Open source software for digital pathology image analysis. Sci Rep. 2017;7(1):16878.

11. Allred DC, Clark GM, Elledge R, Fuqua SA, Brown RW, Chamness GC, et al. Association of p53 protein expression with tumor cell proliferation rate and clinical outcome in node-negative breast cancer. J Natl Cancer Inst. 1993;85(3):200–6.

12. SAS Institute Inc. SAS/STAT ® 14.2 2016

13. Cancer Genome Atlas Research N. The Molecular Taxonomy of Primary Prostate Cancer. Cell. 2015;163(4):1011–25.

14. Richardsen E, Andersen S, Al-Saad S, Rakaee M, Nordby Y, Pedersen MI, et al. Evaluation of the proliferation marker Ki-67 in a large prostatectomy cohort. PLoS One. 2017;12(11):e0186852.

15. Somanathan S, Suchyna TM, Siegel AJ, Berezney R. Targeting of PCNA to sites of DNA replication in the mammalian cell nucleus. J Cell Biochem. 2001;81(1):56–67.

16. Rosenthal SA, Hu C, Sartor O, Gomella LG, Amin MB, Purdy J, et al. Effect of Chemotherapy With Docetaxel With Androgen Suppression and Radiotherapy for Localized High-Risk Prostate Cancer: The Randomized Phase III NRG Oncology RTOG 0521 Trial. J Clin Oncol. 2019;37(14):1159–68.

17. Fei F, Stoddart S, Groffen J, Heisterkamp N. Activity of the Aurora kinase inhibitor VX-680 against Bcr/Abl-positive acute lymphoblastic leukemias. Mol Cancer Ther. 2010;9(5):1318–27.

18. Wu K, Gore C, Yang L, Fazli L, Gleave M, Pong RC, et al. Slug, a unique androgen-regulated transcription factor, coordinates androgen receptor to facilitate castration resistance in prostate cancer. Mol Endocrinol. 2012;26(9):1496–507.

19. Osorio LA, Farfan NM, Castellon EA, Contreras HR. SNAIL transcription factor increases the motility and invasive capacity of prostate cancer cells. Mol Med Rep. 2016;13(1):778–86.

20. Wu K, Zeng J, Zhou J, Fan J, Chen Y, Wang Z, et al. Slug contributes to cadherin switch and malignant progression in muscle-invasive bladder cancer development. Urol Oncol. 2013;31(8):1751–60.

21. Wu ZQ, Li XY, Hu CY, Ford M, Kleer CG, Weiss SJ. Canonical Wnt signaling regulates Slug activity and links epithelial-mesenchymal transition with epigenetic Breast Cancer 1, Early Onset (BRCA1) repression. Proc Natl Acad Sci U S A. 2012;109(41):16654–9.

22. Carpenter RL, Paw I, Dewhirst MW, Lo HW. Akt phosphorylates and activates HSF-1 independent of heat shock, leading to Slug overexpression and epithelial-mesenchymal transition (EMT) of HER2-overexpressing breast cancer cells. Oncogene. 2015;34(5):546–57.

23. Kim D, Langmead B, Salzberg SL. HISAT: a fast spliced aligner with low memory requirements. Nat Methods. 2015;12(4):357–60.

24. Pertea M, Pertea GM, Antonescu CM, Chang TC, Mendell JT, Salzberg SL. StringTie enables improved reconstruction of a transcriptome from RNA-seq reads. Nat Biotechnol. 2015;33(3):290–5.

25. Frazee AC, Pertea G, Jaffe AE, Langmead B, Salzberg SL, Leek JT. Ballgown bridges the gap between transcriptome assembly and expression analysis. Nat Biotechnol. 2015;33(3):243–6.

26. Wang L, Park HJ, Dasari S, Wang S, Kocher JP, Li W. CPAT: Coding-Potential Assessment Tool using an alignment-free logistic regression model. Nucleic Acids Res. 2013;41(6):e74.

27. Georgescu C, Corbin JM, Thibivilliers S, Webb ZD, Zhao YD, Koster J, et al. A TMEFF2-regulated cell cycle derived gene signature is prognostic of recurrence risk in prostate cancer. BMC Cancer. 2019;19(1):423.

28. Chen L, Hong C, Chen EC, Yee SW, Xu L, Almof EU, et al. Genetic and epigenetic regulation of the organic cation transporter 3, SLC22A3. Pharmacogenomics J. 2013;13(2):110–20.

29. Lee Y, Ko D, Min HJ, Kim SB, Ahn HM, Lee Y, et al. TMPRSS4 induces invasion and proliferation of prostate cancer cells through induction of Slug and cyclin D1. Oncotarget. 2016;7(31):50315–32.

30. Vias M, Massie CE, East P, Scott H, Warren A, Zhou Z, et al. Pro-neural transcription factors as cancer markers. BMC Med Genomics. 2008;1:17.

31. Kosaka T, Hongo H, Miyazaki Y, Nishimoto K, Miyajima A, Oya M. Reactive oxygen species induction by cabazitaxel through inhibiting Sestrin-3 in castration resistant prostate cancer. Oncotarget. 2017;8(50):87675–83.

32. Pathak BR, Breed AA, Apte S, Acharya K, Mahale SD. Cysteine-rich secretory protein 3 plays a role in prostate cancer cell invasion and affects expression of PSA and ANXA1. Mol Cell Biochem. 2016;411(1-2):11–21.

33. Grelet S, Andries V, Polette M, Gilles C, Staes K, Martin AP, et al. The human NANOS3 gene contributes to lung tumour invasion by inducing epithelial-mesenchymal transition. J Pathol. 2015;237(1):25–37.

34. Huarte M. The emerging role of lncRNAs in cancer. Nat Med. 2015;21(11):1253–61.

35. Ottman R, Levy J, Grizzle WE, Chakrabarti R. The other face of miR-17-92a cluster, exhibiting tumor suppressor effects in prostate cancer. Oncotarget. 2016;7(45):73739–53.

36. Lu W, Tao X, Fan Y, Tang Y, Xu X, Fan S, et al. LINC00888 promoted tumorigenicity of melanoma via miR-126/CRK signaling axis. Onco Targets Ther. 2018;11:4431–42.

37. Wong RS. Apoptosis in cancer: from pathogenesis to treatment. J Exp Clin Cancer Res. 2011;30:87.

38. Hanahan D, Weinberg RA. Hallmarks of cancer: the next generation. Cell. 2011;144(5):646–74.

39. Medici D, Hay ED, Olsen BR. Snail and Slug promote epithelial-mesenchymal transition through beta-catenin-T-cell factor-4-dependent expression of transforming growth factor-beta3. Mol Biol Cell. 2008;19(11):4875–87.

40. Uygur B, Wu WS. SLUG promotes prostate cancer cell migration and invasion via CXCR4/CXCL12 axis. Mol Cancer. 2011;10:139.

41. Singh S, Sadacharan S, Su S, Belldegrun A, Persad S, Singh G. Overexpression of vimentin: role in the invasive phenotype in an androgen-independent model of prostate cancer. Cancer Res. 2003;63(9):2306–11.

42. Putzke AP, Ventura AP, Bailey AM, Akture C, Opoku-Ansah J, Celiktas M, et al. Metastatic progression of prostate cancer and e-cadherin regulation by zeb1 and SRC family kinases. Am J Pathol. 2011;179(1):400–10.

43. Chen C, Cai Q, He W, Lam TB, Lin J, Zhao Y, et al. AP4 modulated by the PI3K/AKT pathway promotes prostate cancer proliferation and metastasis of prostate cancer via upregulating L-plastin. Cell Death Dis. 2017;8(10):e3060.

44. Murillo-Garzon V, Kypta R. WNT signalling in prostate cancer. Nat Rev Urol. 2017;14(11):683–96.

45. de Aberasturi AL, Calvo A. TMPRSS4: an emerging potential therapeutic target in cancer. Br J Cancer. 2015;112(1):4–8.

46. Kim S, Ko D, Lee Y, Jang S, Lee Y, Lee IY, et al. Anti-cancer activity of the novel 2-hydroxydiarylamide derivatives IMD-0354 and KRT1853 through suppression of cancer cell invasion, proliferation, and survival mediated by TMPRSS4. Sci Rep. 2019;9(1):10003.

47. Kozak J, Wdowiak P, Maciejewski R, Torres A. Interactions between microRNA-200 family and Sestrin proteins in endometrial cancer cell lines and their significance to anoikis. Mol Cell Biochem. 2019;459(1-2):21–34.

48. Shan J, Al-Muftah MA, Al-Kowari MK, Abuaqel SWJ, Al-Rumaihi K, Al-Bozom I, et al. Targeting Wnt/EZH2/microRNA-708 signaling pathway inhibits neuroendocrine differentiation in prostate cancer. Cell Death Discov. 2019;5:139.

49. Chen X, Corbin JM, Tipton GJ, Yang LV, Asch AS, Ruiz-Echevarria MJ. The TMEFF2 tumor suppressor modulates integrin expression, RhoA activation and migration of prostate cancer cells. Biochim Biophys Acta. 2014;1843(6):1216–24.

50. Fu L, Qin YR, Ming XY, Zuo XB, Diao YW, Zhang LY, et al. RNA editing of SLC22A3 drives early tumor invasion and metastasis in familial esophageal cancer. Proc Natl Acad Sci U S A. 2017;114(23):E4631–E40.

