## Supplemental information for "LncRNA *PAINT* is Associated with Aggressive Prostate Cancer and Dysregulation of Slug and Related Genes"

### LncRNA *PAINT* Promotes Prostate Cancer Progression through Modulation of Genes Involved in Epithelial-Mesenchymal Transition and Apoptosis

Md Faqrul Hasan[1§], Kavya Ganapathy[1§], Jiao Sun[2], Ayman, Khatib[1], Thomas Andl[1], Julia N. Saulakova[1], Domenico Coppola[3, 4], Wei Zhang[2] and Ratna Chakrabarti[1*]

**Supplementary Information**

**Cell Proliferation:**

PC-3 cells were transfected with either *PAINT* siRNA smart pool PC-3-*PAINT*$^{si}$ or non-targeting siRNA pool PC-3$^{C}$ (negative control) and incubated for 48h. C4-2B-*PAINT*$^{++}$ stable sublines were seeded and incubated for 48h. Cell proliferation was determined using MTS based Cell Titer Aqueous One Solution Cell Proliferation Assay Kit (Promega) for cell proliferation experiments.

**Flow Cytometry:**

PC-3-*PAINT*$^{si}$, PC-3$^{C}$, C4-2B-*PAINT*$^{++}$ subline and C4-2B$^{C}$ sublines were seeded and harvested at 48h by trypsinization, washed and collected by centrifugation. Cells were permeabilized using absolute methanol for 30min at -20$^{0}$C. Next, PBS was added to the fixed cells and incubated for 5min on ice. Cells were washed with PBS for 3 times, centrifuged and resuspended in PBS with 2% BSA and RNase (100 µg/mL). Cells were incubated for 15min and diluted with PBS containing 2% BSA. Finally, Propidium Iodide (PI) in PBS containing 2% BSA was added and incubated for 30min in dark at room temperature. Fluorescent cells were analyzed in Cytoflex S (Beckman Coulter) Flow Cytometer. Cell cycle analysis was performed using FlowJo software.

**Annexin V Apoptosis Assay**

PC-3-*PAINT*$^{si}$ or PC-3$^{C}$ (negative control) cells were incubated for 48h and treated with DMSO or DTX or VX-680 and incubated for additional 24h. Cells were harvested by trypsinization, washed with PBS, resuspended in annexin V binding buffer and stained with 7-AAD and PE following recommended protocol of PE Annexin V Apoptosis Detection Kit I (BD Biosciences). Cells were incubated in dark for 15min and immediately analyzed in the CytoFLEX S Flow Cytometer (Beckman Coulter Life Sciences). Cells at different quadrants were analyzed using Flow Jo software.

**Migration Assay**

PC-3 cells were seeded in 24 well plates and transfected as previously described. PC-3-*PAINT*$^{si}$ or PC-3$^{C}$

cells were incubated for 48h before making the scratch. C4-2B-*PAINT*$^{++}$ or C4-2B$^{C}$ sublines were seeded and incubated for 48 hours before scratches were made with a 200ul pipette tip and wells were washed 3 times with PBS to remove dislodged cells. Fresh media with 5% FBS was added to each well and images were taken which was considered as 0 hour after making the scratch. To continue with the migration assay, cells were incubated in reduced serum containing media (5% FBS) at 37$^{0}$C in a $CO_2$ incubator and images were taken at 14h (PC-3 cells) and 24h (C4-2B cells) after the scratch was made. Analysis of the rate of cell migration were done using ImageJ analysis software.

**Soft Agar Colony Formation Assay**

C4-2B-*PAINT*$^{++}$ or C4-2B$^{C}$ cells were seeded in semisolid growth medium. Bottom layer of soft agar was generated with 0.5% NuSieve Low melting agar (Lonza) and top layer of soft agar was generated with 0.3% NuSieve Low melting agar (Lonza). Cells were supplemented with 0.5mL complete media every 3 days throughout 2 weeks incubation time. At the end of incubation, cells were washed with PBS and stained with 0.5mL of 0.1% Crystal Violet + 10% ethanol in PBS for 2h. Cells were washed with PBS and incubated overnight at 4$^{0}$C. Next day, cells were washed again with PBS and colonies were visualized in a Leica DMi8 microscope and images (5X) were taken with a Leica DFC450C digital camera attached with the microscope. Colonies were counted and grouped based on their diameter as small (<7 µm) and large (>7 µm). Percentages of small and large colonies were calculated to determine their colony formation properties.

**Immunofluorescence Analysis:**

C4-2B-*PAINT*$^{++}$ or C4-2B$^{C}$ cells were seeded in Chamber slides (Lab-tek Thermo Scientific). After 24h cells were washed with PBS and fixed with 4% paraformaldehyde (PFA) in PBS for 20min on ice. Next, PFA was removed, cells were washed with PBS and permeabilized with 0.1% Triton X-100. Cells were incubated for 30min at room temperature, washed with PBS, and treated with primary antibodies (Ki67) overnight at 4$^{0}$C. Cells were washed with PBS and treated with secondary antibodies for 30min at room temperature. After PBS wash, Phalloidin (25x) staining was done followed by incubation for another 15min at room temperature. Cells were washed in PBS and slides were removed from the chambers and mounted with DAPI Fluoromount-G (Southern Biotech) using coverslips. The prepared slides were imaged using a Leica SP5 confocal microscope and were analyzed with Leica LAS AF software suite. Images were analyzed with Image J software.

**Supplemental Data 1:**

**Table 1:** Patient Criteria in Stages I, II, II and IV

**Supplemental Data 2:**

**Table 2:** Group Sample Sizes for Tumor Tissue Data (n = 53)

**Table 3:** Summary Statistics for Four Primary Measures: Cancer and Normal Tissues. For the cancer tissues (n=54), the patients were about 67 years old, on average (standard deviation is 11.29), the youngest patient was 20 years old and the oldest patient was 82 years old. For the normal tissues (n=3), the patients were about 35 years old, on average (standard deviation is 3.21), the youngest patient was 31 years old and the oldest patient was 37 years old.

**Supplemental Data 3:**

**Table 4:** Summary Statistics for Primary Cancer Tissue Measures by Stage, Grade, GSI and Metastasis Indicator: Cancer Tissue Data (n= 53)

**Supplemental Data 4:**

**Table 5:** Analysis of Maximum Likelihood Estimates

**Supplemental Data 5:**

**Table 6:** Sequencing statistics of each sample used for RNA-seq

**Figure S1.** A: Survival analysis based on TCGA PRAD dataset showing prostate cancer patients with higher *PAINT* expression (N = 149) have lower survival rate compared to patients with lower *PAINT* expression (N = 148). **p-value* = 0.07. B: Bright field images of PC-3 cell morphology upon inhibition of *PAINT* showing more epithelial-like cell shape compared to the elongated shape of the control siRNA transfected cells. C: Analysis of Ki67 positive cells in C4-2B-$PAINT^{++}$ and C4-2B$^{C}$ cells. Data show mean ± SD of at least three biological replicates.

**Figure S2**. A: The heatmap for the relations among samples showing the Pearson correlation of gene level expression of all samples above 0.996. B: 3D scatter plot shows novel coding potential distribution of coding transcripts (red dots) and non-coding transcripts (green dots). Logistic regression model used to determine the protein coding potential used four features (ORF size and coverage, Hexamer score and Fickett score). C: Circular plot showing the relationship between 18 downregulated genes in C4-2B-$PAINT^{++}$ cells and various biological processes. D: Circular plot showing the distribution of 20 downregulated genes in C4-2B-$PAINT^{++}$ cells with relation to cellular components. E: Circular plot of GO enrichment analysis showing the relationship between 20 upregulated genes in C4-2B-$PAINT^{++}$ cells and various biological processes. F: Circular plot of GO enrichment analysis showing the distribution of 20 upregulated genes in C4-2B-$PAINT^{++}$ with relation to cellular components.

**Figure S3**. A-D: Top 4 downregulated KEGG pathways showing specific genes that were dysregulated in mucin type O-Glycan biosynthesis (A), aldosterone regulated sodium reabsorption (B), endocrine and other factor regulated calcium reabsorption (C), and cGMP-PKG signaling pathway(D) in C4-2B-$PAINT^{++}$ cells. (G-I) Top 3 upregulated KEGG pathways showing specific genes that were dysregulated in p53 signaling pathway (G), Alanine aspartate and glutamate metabolism (H) Arachidonic acid metabolism (I) in C4-2B-$PAINT^{++}$ cells.

**Table 1. Patient Criteria in Stages I, II, II and IV**

| No | Grade | Stage | Gleason Scores | TNM | No | Grade | Stage | Gleason Scores | TNM | No | Grade | Stage | Gleason Scores | TNM |
|---|---|---|---|---|---|---|---|---|---|---|---|---|---|---|
| 1 | 1 | I | 2+2 | T1N0M0 | 18 | 2 | II | 3+3 | T2N0M0 | 5 | 2 | III | 2+4 | T3N0M0 |
| 1 | 1 | II | 1+2 | T2N0M0 | 19 | 3 | II | 4+5 | T2N0M0 | 6 | 2 | III | 2+4 | T3N0M0 |
| 2 | 1 | II | 1+2 | T2N0M0 | 20 | 3 | II | 4+5 | T2N0M0 | 7 | 2 | III | 3+4 | T3N0M0 |
| 3 | 1 | II | 2+2 | T2aN0M0 | 21 | 2 | II | 3+5 | T2N0M0 | 8 | 2--3 | III | 4+3 | T3aN0M0 |
| 4 | 1 | II | 1+2 | T2N0M0 | 22 | 3 | II | 4+4 | T2N0M0 | 9 | 2 | III | 3+3 | T3N0M0 |
| 5 | 2 | II | 2+3 | T2N0M0 | 23 | 3 | II | 4+4 | T2N0M0 | 10 | 3 | III | 5+5 | T3N0M0 |
| 6 | 2 | II | 2+4 | T2N0M0 | 24 | 3 | II | 5+4 | T2N0M0 | 11 | 3 | III | 4+5 | T3N0M0 |
| 7 | 1 | II | 1+2 | T2N0M0 | 25 | 3 | II | 5+4 | T2N0M0 | 1 | 1 | IV | 1+2 | T3N1M1 |
| 8 | 2 | II | 3+3 | T2N0M0 | 26 | 3 | II | 5+5 | T2N0M0 | 2 | 1 | IV | 2+2 | T4N1M1 |
| 9 | 2 | II | 3+3 | T2aN0M0 | 27 | 3 | II | 5+4 | T2N0M0 | 3 | 2 | IV | 2+4 | T3N2M1 |
| 10 | 2 | II | 3+3 | T2N0M0 | 28 | 3 | II | 5+5 | T2aN0M0 | 4 | 2 | IV | 3+3 | T2N1M1c |
| 11 | 2 | II | 3+3 | T2N0M0 | 29 | 3 | II | 4+4 | T2N0M0 | 5 | 2 | IV | 2+4 | T3N1M0 |
| 12 | 2 | II | 2+4 | T2N0M0 | 30 | 3 | II | 5+4 | T2N0M0 | 6 | 2 | IV | 3+3 | T3N1M0 |
| 13 | 2 | II | 3+3 | T2N0M0 | 31 | 3 | II | 5+5 | T3N0M0 | 7 | 2 | IV | 3+4 | T4N1M1b |
| 14 | 2 | II | 2+4 | T2N0M0 | 1 | 1 | III | 2+2 | T3N0M0 | 8 | 3 | IV | 4+5 | T3N1M0 |
| 15 | 2 | II | 3+3 | T2N0M0 | 2 | 1 | III | 2+2 | T3N0M0 | 9 | 3 | IV | 5+4 | T4N0M0 |
| 16 | 3 | II | 4+4 | T2N0M0 | 3 | 3 | III | 4+5 | T3N0M0 | 10 | 3 | IV | 5+5 | T3N1M0 |
| 17 | 3 | II | 4+4 | T2N0M0 | 4 | 2--3 | III | 4+3 | T3N0M0 | 11 | 3 | IV | 5+4 | T3N1M0 |

**Table 2. Group Sample Sizes for Tumor Tissue Data (n = 53)**

| Measure | Group Sample Size | Percentage (%) |
|---|---|---|
| Stage 3 | | |
| I/II | 31 | 58.49 |
| III | 11 | 20.75 |
| IV | 11 | 20.75 |
| Grade | | |
| 1 | 10 | 18.87 |
| 2 | 21 | 39.62 |
| 3 | 22 | 41.51 |
| Gleason Scores Indicator | | |
| Low (6 or less) | 28 | 52.83 |
| High (7 or grater) | 25 | 47.17 |
| Metastasis | | |
| Absent | 43 | 81.13 |
| Present | 10 | 18.87 |

**Table 3. Summary Statistics for Four Primary Measures: Cancer and Normal Tissues**

| **Measure** | **Mean** | **Standard Deviation** | **Minimum** | **Maximum** |
|---|---|---|---|---|
| **Cancer Tissue (n=53)** | | | | |
| Target 1 | 8.59 | 8.1181 | 0.95 | 45.81 |
| Target 2 | 7.29 | 6.2073 | 0.51 | 25.21 |
| Target 3 | 8.07 | 5.9992 | 1.58 | 23.61 |
| Average | 7.98 | 6.3347 | 1.44 | 30.22 |
| **Normal Tissue (n=3)** | | | | |
| Target 1 | 3.65 | 0.5688 | 3.00 | 4.07 |
| Target 2 | 3.93 | 0.5225 | 3.41 | 4.45 |
| Target 3 | 3.44 | 1.4411 | 2.28 | 5.05 |
| Average | 3.67 | 0.6051 | 3.19 | 4.35 |

Target 1, 2 and 3 are staining of the TMA slide in triplicate

**Table 4. Summary Statistics for Primary Cancer Tissue Measures by Stage, Grade, GSI and Metastasis Indicator: Cancer Tissue Data (n= 53)**

| Group | Measure | Mean | Standard Deviation | Median |
|---|---|---|---|---|
| **Stage** | | | | |
| I/II (n=31) | Target 1 | 6.29 | 4.49 | 5.30 |
| | Target 2 | 5.37 | 3.60 | 4.97 |
| | Target 3 | 6.34 | 3.73 | 5.18 |
| | Average | 6.00 | 3.68 | 5.10 |
| III (n=11) | Target 1 | 8.16 | 7.81 | 3.94 |
| | Target 2 | 6.54 | 6.89 | 3.50 |
| | Target 3 | 7.59 | 6.87 | 4.01 |
| | Average | 7.43 | 6.91 | 3.43 |
| IV (n=11) | Target 1 | 15.51 | 12.34 | 12.55 |
| | Target 2 | 13.44 | 7.77 | 13.11 |
| | Target 3 | 13.45 | 7.53 | 13.82 |
| | Average | 14.13 | 8.14 | 14.19 |
| **Grade** | | | | |
| 1 (n=10) | Target 1 | 11.32 | 7.93 | 11.26 |
| | Target 2 | 9.84 | 6.53 | 8.07 |
| | Target 3 | 10.55 | 7.96 | 9.24 |
| | Average | 10.57 | 7.35 | 9.43 |
| 2 (n=21) | Target 1 | 5.98 | 4.41 | 6.65 |
| | Target 2 | 6.54 | 5.74 | 3.97 |
| | Target 3 | 6.61 | 4.10 | 5.92 |
| | Average | 6.38 | 4.50 | 5.46 |
| 3 (n=22) | Target 1 | 9.85 | 10.28 | 6.24 |
| | Target 2 | 6.84 | 6.47 | 4.79 |
| | Target 3 | 8.34 | 6.39 | 6.15 |
| | Average | 8.35 | 7.14 | 5.96 |

| Group | Measure | Mean | Standard Deviation | Median |
|---|---|---|---|---|
| **GSI** | | | | |
| Low (n=28) | Target 1 | 7.94 | 6.25 | 7.25 |
| | Target 2 | 7.84 | 6.11 | 5.59 |
| | Target 3 | 8.14 | 5.89 | 6.51 |
| | Average | 7.98 | 5.84 | 6.40 |
| High (n=25) | Target 1 | 9.32 | 9.89 | 5.32 |
| | Target 2 | 6.66 | 6.38 | 4.62 |
| | Target 3 | 8.00 | 6.24 | 5.27 |
| | Average | 7.99 | 6.97 | 5.85 |
| **Metastasis** | | | | |
| Absent (n=43) | Target 1 | 7.69 | 8.06 | 5.30 |
| | Target 2 | 6.09 | 5.31 | 4.45 |
| | Target 3 | 7.01 | 5.15 | 5.18 |
| | Average | 6.93 | 5.89 | 5.10 |
| Present (n=10) | Target 1 | 12.48 | 7.54 | 11.45 |
| | Target 2 | 12.43 | 7.39 | 12.73 |
| | Target 3 | 12.66 | 7.45 | 12.67 |
| | Average | 12.52 | 6.48 | 13.39 |

**Table 5. Analysis of Maximum Likelihood Estimates**

| Parameter | | DF | Estimate | Standard Error | Wald Chi-Square | Pr > ChiSq |
|---|---|---|---|---|---|---|
| Intercept | | 1 | -6.3207 | 4.6017 | 1.8867 | 0.1696 |
| Age | | 1 | 0.0351 | 0.0637 | 0.3030 | 0.5820 |
| Average | | 1 | 0.2926 | 0.0979 | 8.9376 | 0.0028 |
| Grade | 2 | 1 | 0.7795 | 0.6574 | 1.4060 | 0.2357 |
| Grade | 3 | 1 | -0.5172 | 0.6834 | 0.5728 | 0.4492 |

**Table 6. Sequencing Statistics of Samples**

| Sample Name | Total Read Counts | Raw pairs | Trimmed Reads | Mapped Reads | Identified Gene Counts | Identified Transcript counts |
|---|---|---|---|---|---|---|
| C1 | 54745196 | 27372598 | 27365149 | 92.73% | 11575 | 27276 |
| C2 | 56299252 | 28149626 | 28147099 | 92.87% | | |
| C3 | 53941034 | 26970517 | 26963264 | 92.60% | | |
| E1 | 69409542 | 34704771 | 34701416 | 93.92% | 11739 | 27554 |
| E2 | 58151332 | 29075666 | 29075494 | 92.49% | | |
| E3 | 63337080 | 31668540 | 31662191 | 93.77% | | |

**Raw pairs** → Raw sequencing fragment numbers
**Trimmed Reads** → Fragments number (read pairs) after 5’, 3’-adaptor trimmed and filtered 20 bp reads.
**Mapped %** → The proportion of Reads number aligning to reference genome with Hisat 2 software in Trimmed pairs.

**The number of identified genes and transcripts per group was calculated based on the mean of FPKM in group >= 0.5.**

Supplemental Data 5

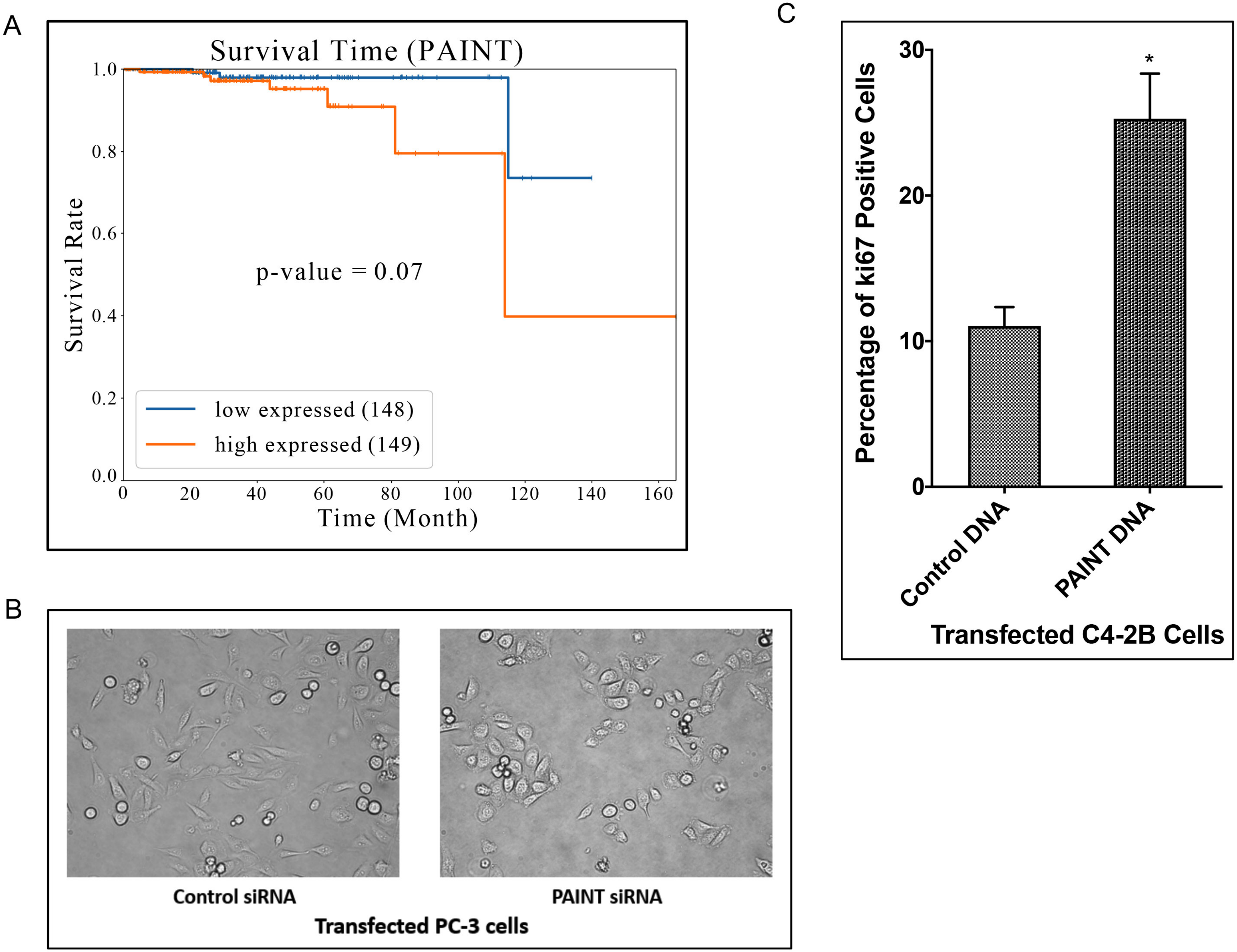
A
Survival Time (PAINT)
Survival Rate
1.0
0.8
0.6
0.4
0.2
0.0
0
20
40
60
80
100
120
140
160
Time (Month)
p-value = 0.07
low expressed (148)
high expressed (149)
B
Control siRNA
PAINT siRNA
Transfected PC-3 cells
C
Percentage of ki67 Positive Cells
30
20
10
0
*
Control DNA
PAINT DNA
Transfected C4-2B Cells

Fig S1

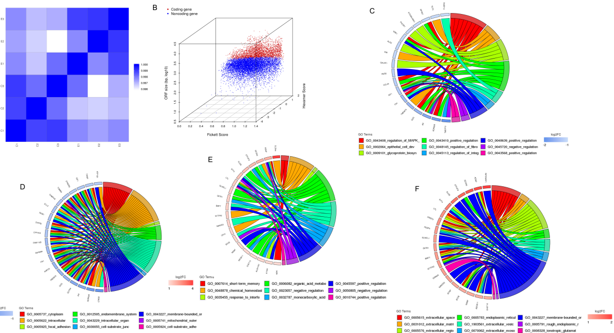

B
Coding gene
Noncoding gene
ORF size (bp, log10)
Fickett Score
Hexamer Score
C
GO Terms
GO_0043408_regulation_of_MAPK_
GO_0002064_epithelial_cell_dev
GO_0009101_glycoprotein_biosyn
GO_0043410_positive_regulation
GO_0048145_regulation_of_fibro
GO_0045113_regulation_of_integ
GO_0048639_positive_regulation
GO_0045720_negative_regulation
GO_0043568_positive_regulation
log2FC
D
log2FC
GO Terms
GO_0005737_cytoplasm
GO_0005622_intracellular
GO_0005925_focal_adhesion
GO_0012505_endomembrane_system
GO_0043229_intracellular_organ
GO_0030055_cell-substrate_junc
GO_0043227_membrane-bounded_or
GO_0005741_mitochondrial_outer
GO_0005924_cell-substrate_adhe
E
log2FC
GO Terms
GO_0007614_short-term_memory
GO_0048878_chemical_homeostasi
GO_0035455_response_to_interfe
GO_0006082_organic_acid_metabo
GO_0023057_negative_regulation
GO_0032787_monocarboxylic_acid
GO_0045597_positive_regulation
GO_0050805_negative_regulation
GO_0010744_positive_regulation
F
GO Terms
GO_0005615_extracellular_space
GO_0031012_extracellular_matri
GO_0005576_extracellular_regio
GO_0005783_endoplasmic_reticul
GO_1903561_extracellular_vesic
GO_0070062_extracellular_exoso
GO_0043227_membrane-bounded_or
GO_0005791_rough_endoplasmic_r
GO_0008328_ionotropic_glutamat
log2FC


Fig S2

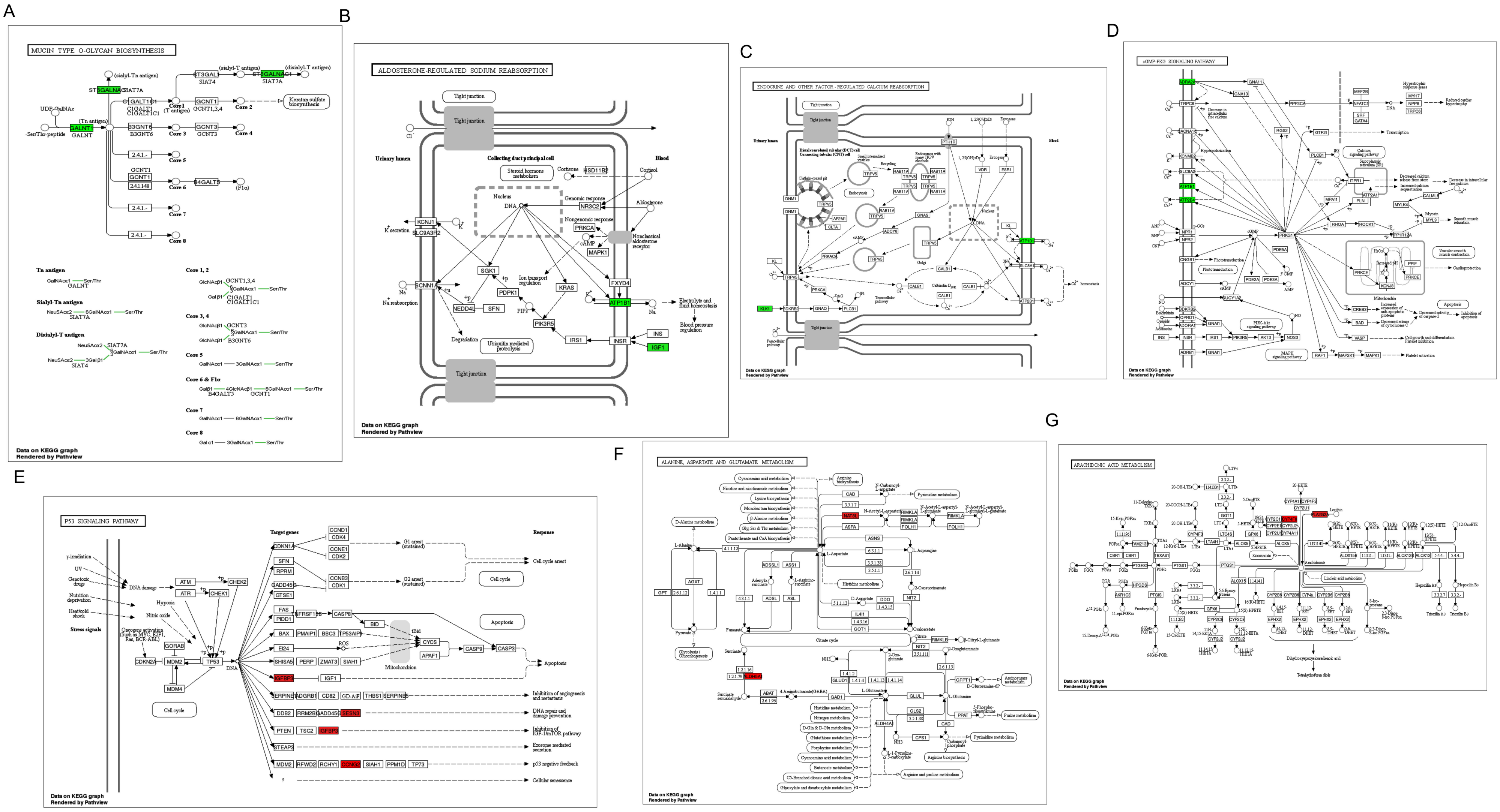
A
MUCIN TYPE O-GLYCAN BIOSYNTHESIS
B
ALDOSTERONE-REGULATED SODIUM REABSORPTION
C
ENDOCRINE AND OTHER FACTOR-REGULATED CALCIUM REABSORPTION
D
cGMP-PKG SIGNALING PATHWAY
E
P53 SIGNALING PATHWAY
F
ALANINE, ASPARTATE AND GLUTAMATE METABOLISM
G
ARACHIDONIC ACID METABOLISM
Data on KEGG graph
Rendered by Pathview

Fig S3
